# Discovery of Drug-like Ligands for the Mac1 Domain of SARS-CoV-2 Nsp3

**DOI:** 10.1101/2020.07.06.190413

**Authors:** Rajdeep S. Virdi, Robert V. Bavisotto, Nicholas C. Hopper, Nemanja Vuksanovic, Trevor R. Melkonian, Nicholas R. Silvaggi, David N. Frick

## Abstract

Small molecules that bind the SARS-CoV-2 non-structural protein 3 Mac1 domain in place of ADP-ribose could be useful as molecular probes or scaffolds for COVID-19 antiviral drug discovery because Mac1 has been linked to coronavirus’ ability to evade cellular detection. A high-throughput assay based on differential scanning fluorimetry (DSF) was therefore optimized and used to identify possible Mac1 ligands in small libraries of drugs and drug-like compounds. Numerous promising compounds included nucleotides, steroids, beta-lactams, and benzimidazoles. The main drawback to this approach was that a high percentage of compounds in some libraries were found to influence the observed Mac1 melting temperature. To prioritize DSF screening hits, the shapes of the observed melting curves and initial assay fluorescence were examined, and the results were compared with virtual screens performed using Autodock VINA. The molecular basis for alternate ligand binding was also examined by determining a structure of one of the hits, cyclic adenosine monophosphate, with atomic resolution.

## Introduction

Direct-acting antivirals (DAAs) are desperately needed to treat COVID-19 patients and stem the devastation caused by the current SARS-CoV-2 pandemic. DAAs are typically developed from potent inhibitors of viral enzymes or high-affinity ligands of viral proteins. For example, remdesivir inhibits the SARS-CoV-2 RNA dependent RNA polymerase, halting SARS-CoV-2 replication.^1^ Based on past experiences, any effective DAA therapy will likely require a cocktail of more than one antiviral agent because drug resistance evolves rapidly. Methods are therefore needed to rapidly identify small-molecule drug-like ligands for as many SARS-CoV-2 proteins as possible. The ∼29,900 nucleotide SARS-CoV-2 genome encodes many potential DAA targets including 16 nonstructural proteins (nsps), four structural proteins, six accessory proteins, and possibly many others. Most SARS-CoV-2 nsps are products of the rep1b open-reading frame that encodes a short (*aka* ORF1a) and long (*aka* ORF1b) polyprotein because an internal RNA hairpin occasionally causes translational frameshifting. The subject of this study is the multifunctional 945 amino-acid-long nsp3 protein that cleaves the three junctures separating nsp1, nsp2 and nsp3.^2^ SARS-CoV-2 nsp3 is most likely tethered to the ER with two ubiquitin-like domains (Ubl1 and Ubl2), two papain-like protease domains (PLP1^pro^ and PLP2^pro^), three macrodomains (Mac1, Mac2, and Mac3), a nucleic-acid-binding domain, and a hypervariable region facing the cytoplasm.

In a previous study, Frick *et al*.^3^ characterized the SARS-CoV-2 Mac1 domain (*aka* the “X” domain) and its ability to binds ADP-ribose (ADPr). Here, we report the results of pilot screens designed to find drug-like Mac1 domain ligands, which might facilitate DAA design, or which could be useful as molecular probes. A differential scanning fluorimetry (DSF, aka the thermal shift, or ThermoFluor)^4–6^ assay was optimized and used to screen 726 compounds in the National Institutes of Health clinical collection (NIHcc), the National Cancer Institute (NCI) mechanistic set (540 compounds), and Sigma-Aldrich’s 1280 compound Library of Pharmacologically Active Compounds (LOPAC^1280^). Since up to 5% of compounds in each set influenced the apparent melting temperature, compounds were prioritized using information derived from individual melting curves and virtual screens performed with AutoDock VINA.^7^ A high-resolution structure of one hit compound, cyclic adenosine monophosphate (cAMP) was determined in complex with SARS-CoV-2 Mac1, revealing at atomic resolution the capacity of the binding cleft to accommodate other ligands.

## Materials and Methods

Purified SARS-CoV-2 Mac1 protein was prepared as described previously. ^3^ DSF Assays were performed in 96-well PCR plates using an Eppendorf Mastercycler ep Realplex Quantitative Realtime PCR System with each well containing 19 µl of Master-mix (5 µM Mac1) and 1 µl of a compound stock (10 mM for screening) or DMSO. The master-mix was prepared by adding 20 µl of 500 µM Mac1, 2.5 µl 5000x SPYRO Orange protein gel stain (Sigma-Aldrich catalog #S5692) to 1977.5 µl of buffer (20 mM MOPS, 25 mM NaCl, pH 7). The 96-well PCR plate was then sealed by a clear adhesive film and centrifuged at 1,100 RPM for 5 mins. The temperature was raised from 20 °C to 95 °C at a rate of 2 °C per min while measuring the fluorescence in the “TAMRA” channel. Each plate included both negative (DMSO) and positive (ADPr) controls. T_m_’s were calculated by fitting the data to equation 1 using either GraphPad Prism or TSA-CRAFT (https://sourceforge.net/projects/tsa-craft/).^8^

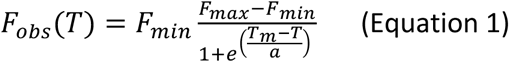

In Eq. 1, F_obs_ (T) is the observed fluorescence at each temperature (T), F_min_ is the minimum observed fluorescence, F_max_ is maximum observed fluorescence, and a is a *Hill* slope. Two methods were used estimate the affinity of Mac1 from DSF. First, the observed melting temperatures were plotted versus ligand and fit to Eq. 2 to determine the amount of compound needed to cause a change the melting temperature of 50% (EC_50_) using equation 2, and a non-linear regression to estimate ΔT_m max_ (the maximum change in T_m_) from the melting temperature of Mac1 in the absence of ligand (T_m 0_)

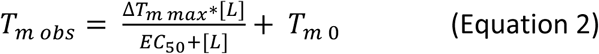

The dissociation constant (K_d_) of Mac1 and ADPr and the equilibrium constant describing protein unfolding (K_u_) were also estimated using isothermal analysis as described by Bai et al.^9^ Briefly, normalized melting curves were used to calculate the fraction of protein unfolded at a particular temperature (*f*_u_), and those values fitted to the total ligand (L_t_) and protein (P_t_) concentrations using non-linear regression and Eq. 3:

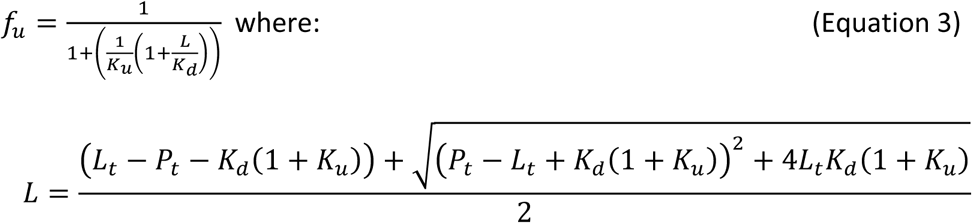

Computational ligand screening was performed using both the unligated (6WEY and 6VXS) and ADPr-bound forms (6W02), using the program AutoDock VINA.^7^ The protein files were downloaded directly from the protein data bank and processed as described below before submitting for screening. All solvent molecules (HETATM) were removed from the files. Polar hydrogen atoms were added and Kollman charges were included in the protein files. The converted protein and ligand file pdbqt libraries were uploaded to a parallel computing cluster and run with the following parameters: Energy difference = 4; Number of recorded modes = 20; Exhaustiveness was set to 12. Docking box location was configured prior to using AutoDock tools. After the docking calculation was complete, the locations, orientations, and binding affinities of the top candidates were examined using UCSF Chimera and tabulated for comparison.

Mac1 was prepared for crystallization as described before^3^ with the following modifications. First, the plasmid was modified to express one additional N-terminal residue (E206) and C-terminus was shortened by three residues such that it encoded residues 206-374 of the SARS-Cov-12 nsp3. After purification and TEV protease cleavage the tag-free protein was concentrated to 20 mg/ml in 20 mM HEPES, pH 7.5, 150 mM NaCl and 1 mM TCEP. Crystallization was accomplished as described for the Mac1·AMP complex^10^ (1 µl of concentrated Mac1 was mixed with 1 µl 30% PEG 4000, 0.1 M MES, pH 6.5). Plate-shaped crystals grew in 3-7 days at 22 °C.

The Mac1·cAMP complex was prepared by soaking the crystal in a solution containing 35% PEG 4000 and 20 mM cAMP for 30 min. Cryo-protection was accomplished by briefly soaking the crystal in 35% PEG 4000, 20 mM cAMP, and 20% glycerol before plunging it into liquid nitrogen. Diffraction data were collected on Life Sciences Collaborative Access Team (LS-CAT) beamline 21-ID-F at the Advanced Photon Source of Argonne National Laboratory, which is fitted with a fixed-wavelength beam at 0.97872 Å and a MarMosaic M300 detector. The data were collected with an oscillation width of 0.5° per image for a total oscillation of 180°. The data were processed using HKL2000;^11^ data collection statistics are provided in Table S1.

The structure was determined by molecular replacement in PHASER^12^ using PDB ID 6WEY,^3^ with solvent molecules and B-factor information removed, as the search model. The model underwent iterative rounds of (re)building in COOT^13^ and refinement in PHENIX.refine.^14, 15^ Translation-libration-screw (TLS) refinement provided a more realistic treatment of the atomic displacement parameters; TLS groups were identified by phenix.find_tls_groups. Model refinement and validation statistics are provided in Table S1. The coordinates were deposited in the PDB (accession code 7JME).

## Results

### An optimized DSF assay for SARS-CoV-2 Mac1

DSF has been used previously to study ligand binding to viral macrodomains.^16,17^ In DSF experiments using SARS-CoV-2 Mac1, the presence of ADPr raised the Mac1 melting temperature in a concentration-dependent manner (Fig. 1A). To estimate the ligand concentrations needed to alter melting temperatures by 50% (EC_50_), melting temperatures were fit to Eq. 2 (Fig. 1B). Such EC_50_ values do not, however, describe protein-ligand affinity because DSF assays do not directly measure binding. The isothermal analysis recently described by Bai *et al*.^9^ was therefore used to estimate binding affinities. Fits of the fraction of protein unfolded at various temperatures in the presence of various ligand concentrations (Fig. 1C) were used to estimate a dissociation constant (K_d_) and equilibrium unfolding constant (K_u_). Each depended on temperature (Fig. 1D), and when fit to the Van’t Hoff relationship, were in good agreement with the dissociation constant describing the interaction of ADPr and Mac1 (10 µM) determined at 23 °C.^3^

**Figure 1.**
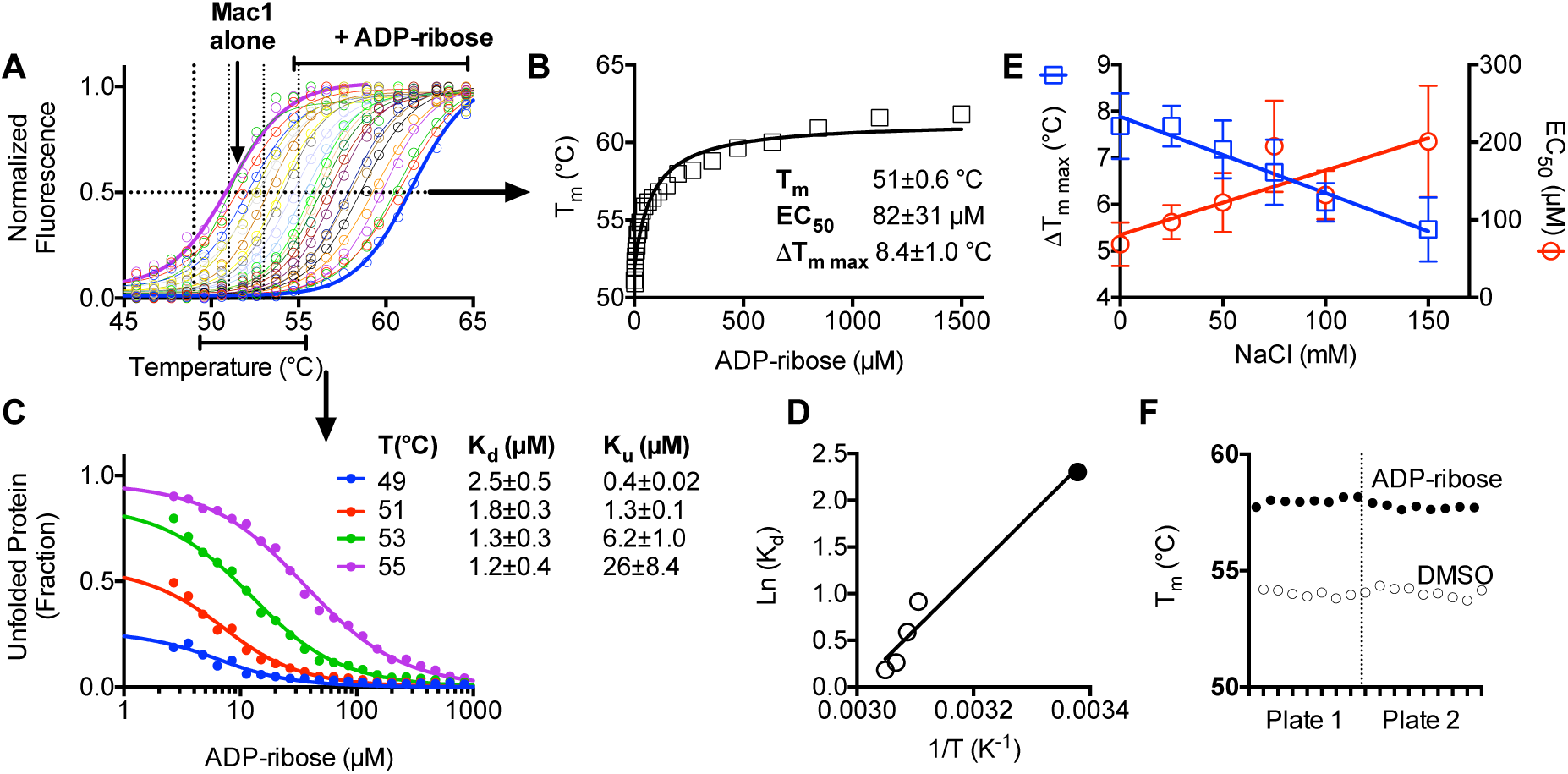
DSF assay optimization. (A) Normalized SPYRO Orange fluorescence in the presence of 6 µM Mac1 protein at various temperatures in the presence of indicated concentrations of ADPr. Data are fit to Eq. 1 using non-linear regression with GraphPad Prism. (B) T_m_ values obtained from direct fitting to Eq. 1. Data are fit to Eq. 2. (B) Isothermal analysis^9^ of percent unfolded protein at each indicated temperature. Data are fit to Eq. 3 with indicated constants. Uncertainties are standard errors of the curve fits. (C) Van’t hoff plot of estimated K_d_ values from *panel C* (open circles) and the K_d_ for ADPr binding to Mac1 that was previously determined at 23 °C using isothermal titration calorimetry (filled circle).^3^ (E) ADPr titrations in 20 mM MOPS pH buffer supplemented with indicated NaCl concentrations. Plotted are best-fit ΔT_m max_ (squares, left y-axis) and EC_50_ (circles, right axis). Error bars mark standard errors in the curve fits. (F) T_m_ values for positive (500 µM ADPr) and negative controls (DMSO only) from two different 96-well plates (Z’ factor = 0.72).

Dimethyl sulfoxide (DMSO) did not change the melting curve even at concentrations as high as 10% (v/v) and similar results were also obtained when titrations with ADPr were repeated in various buffers, with the pH ranging from 6.5 to 8.0, or in the presence of various concentrations of divalent metal cations (Mg^2+^ or Mn^2+^). In contrast, the ionic strengths of the assay buffers influenced the results, with the largest ΔT_m_ values and lowest EC_50_ values being obtained at the lowest ionic strengths (Fig. 1E). Based on these results, DSF assays were subsequently performed in 20 mM MOPS pH 7 buffer containing 25 mM NaCl to reduce possible non-specific interactions with ligands. Z’ factors^18^ were always above 0.5 for each plate and typically above 0.7. Plate-to-plate variability was negligible (Fig. 1F). The first screen was performed using compounds from an NCI library (NCI, https://dtp.cancer.gov/repositories/)(Fig. 2A),^19^ the second was performed using the NIHcc (https://commonfund.nih.gov/molecularlibraries/tools) (Fig. 2B),^20^ and the third using Sigma-Aldrich’s LOPAC^1280^ (Fig. 2C, D). Although ligands that reduce a protein’s T_m_ are often assumed to bind and stabilize unfolded structures,^21^we nevertheless also examined some of these hits in more detail. In addition to nucleotides suspected to bind in place of ADPr, other noteworthy hits relevant to current COVID-19 research included the ACE inhibitor telmisartan, several steroids, and β-lactam antibiotics (Fig. 2E).

**Figure 2.**
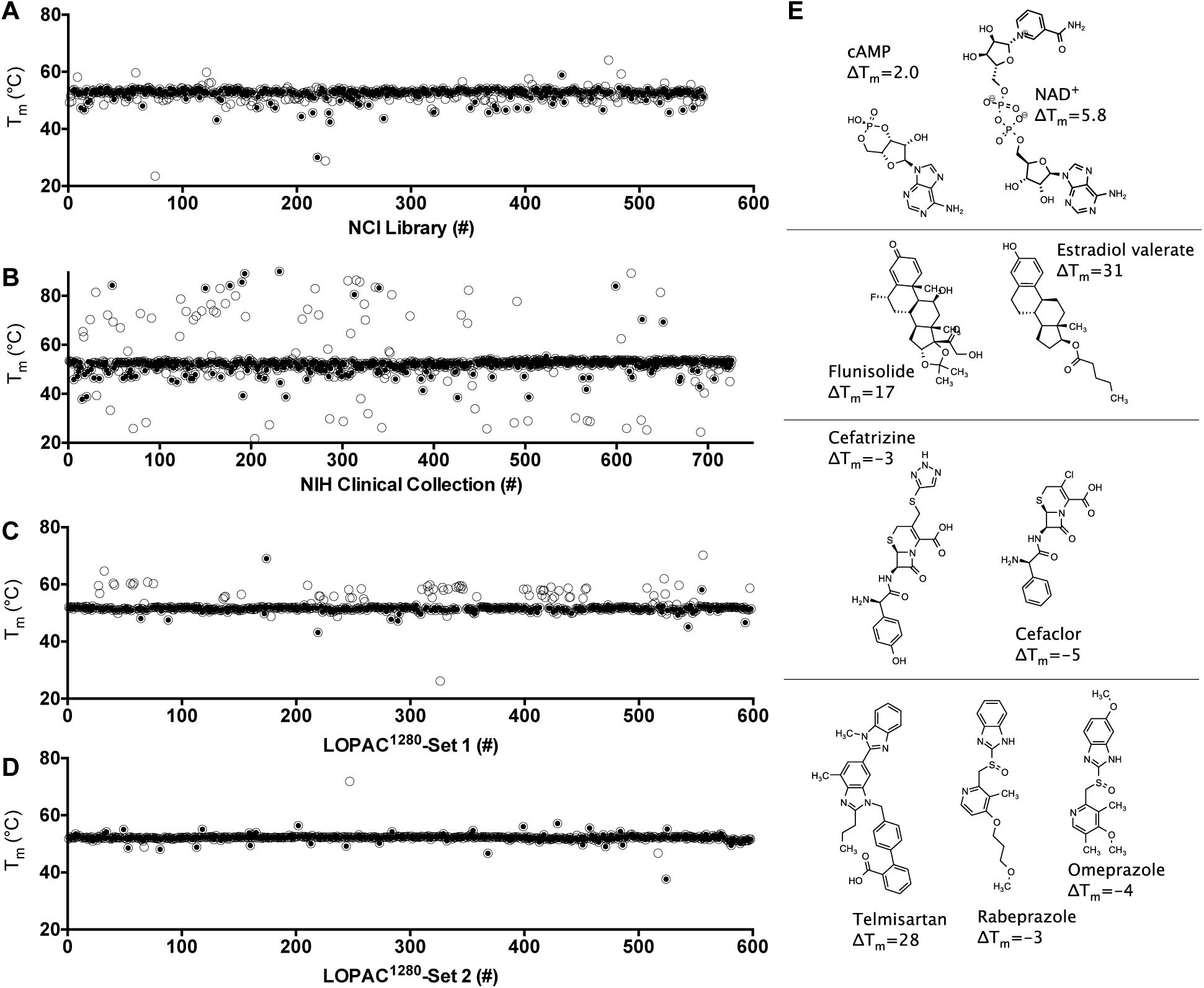
DSF screens of FDA-approved drugs and drug-like compounds for SARS-CoV-2 Mac1 ligands. T_m_ values calculated by fitting melting curves to Eq. 1 (open circles) obtained for Mac1 in the presence of each compound in the NCI library (A) the NIHcc (B) and the LOPAC^1280^ (C,D). Assays yielding a “typical” melting curve, as defined by the TSA-CRAFT algorithm, are noted (filled circles). (E) Selected hit compounds separated based chemotype: nucleotides, steroids, β-lactam antibiotics, and benzimidazoles.

### Prioritizing hits in DSF screens

Screening results reveal that the major limitation of this approach is that a high percentage of compounds in some libraries influence the observed protein melting temperature, which was particularly evident with the NIHcc. Many of these compounds either quench fluorescence, fluoresce themselves, or interact with the reporter dye. To exclude such compounds in follow-up experiments, the TSA-CRAFT software package was used to identify what it defines as “typical” curves (filled-circles, Figs. 2, 3). We found that such interfering compounds could also be identified by simply plotting T_m_ values versus the initial fluorescence seen in the melting curve (Fig. 3A).

**Figure 3.**
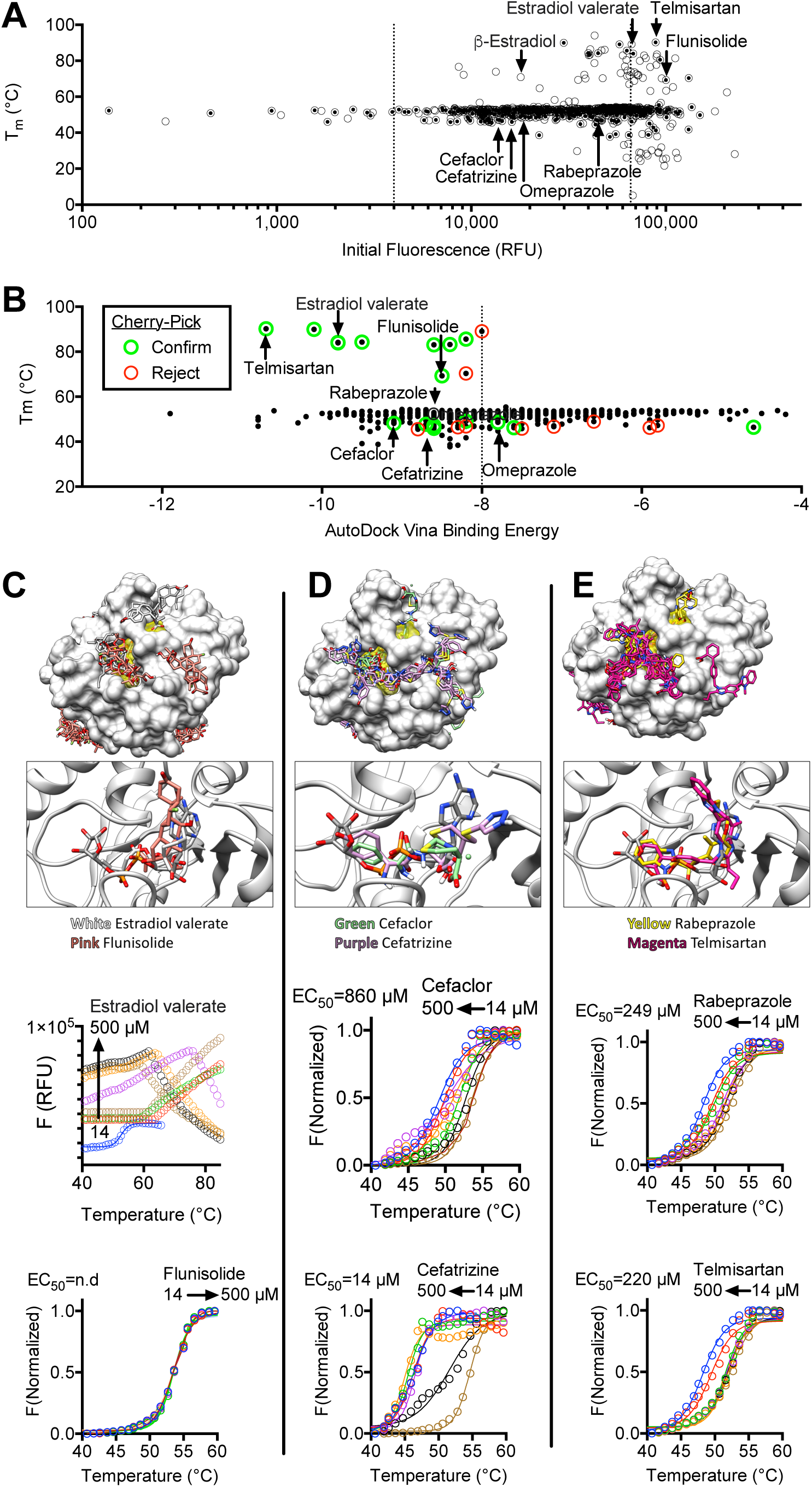
Methods to prioritize hits from DSF screens. (A) Plot of T_m_ values for samples in the NIHcc plotted versus fluorescence observed at the beginning of each melt (*i*.*e*. @ 20 °C). “Typical” melting curves are highlighted (filled circles). The dotted lines are arbitrary cutoffs drawn at three times more and less than the average fluorescence intensity recorded in all assays. (B) AutoDock Vina binding energies obtained for each compound after docking with PDB file 6WEY (y-axis) compared with the T_m_ derived using TSA-CRAFT. (C-E) Representative Structures obtained using AutoDock Vina Virtual Screening. *Top panels* show the top 20 binding modes for selected compounds, and the bottom panels show the top binding mode for each compound compared to the ADPr bound in PDB file 6W02. The ADPr binding cleft is highlighted in yellow. Concentration response analysis of each compound is DSF assays are shown, along with EC_50_ values.

As another method for hit prioritization, “virtual” screens were performed with SARS-CoV-2 Mac1 crystal structures (PDB files 6WEY, 6W02, and 6VXS) as targets in the program AutoDock Vina (Fig. 3A). Each was searched free from ligands. Binding sites were not restricted but, for most compounds, minimum binding energy (best fit) values were obtained for structures in which the compound docked near the ADPr binding site. Plots of AutoDock Vina scores *vs*. T_m_ could be used to identify compounds for follow-up assays (Fig. 3B). First, there was a clear correlation between the number of hits and AutoDock Vina scores, with more hits clustering at lower energies. Second, when cherry-pick assays were performed on hits, those with lower energy scores (12/15) were more likely to be reproducible than compounds with higher-energy scores (3/9) (Fig. 3B). Close examination of molecular models generated by AutoDock Vina reveal that the steroids (Fig. 3C), β-lactams (Fig. 3D) and benzimidazoles (Fig. 3E) could each occupy the ADPr-binding cleft on SARS-CoV-2 Mac1. The larger compounds in each class make more contacts with amino acids in the cleft, explaining their higher binding energies.

By combining these prioritization methods, compounds that more likely bind Mac1 could be differentiated from those that likely do not. For example, all nucleotides that were hits yielded results similar to those seen with ADPr. However, the steroids either yielded atypical curves (β-estradiol) or increased the DSF assay fluorescence (estradiol valerate and flunisolide). When new aliquots of selected compounds were purchased, abnormal melting curves were observed with estradiol valerate and no effects were observed with flunisolide (Fig. 3C). In contrast, fresh batches of both lactams and two benzimidazoles yielded the same effects seen with the screening library (Fig. 3D, E). Interestingly, fresh telmisartan yielded a different effect, lowering the apparent T_m_ as was seen with related compounds (Fig. 3D), suggesting that a possible degradation product led to the T_m_ increase observed using the library sample.

### Structure of cAMP bound to SARS-CoV-2 Mac1

The protein construct used to determine the structure of the apoenzyme (PDB ID 6WEY; nsp3 residues 207-377) crystallized with such tight packing that it proved impossible to obtain structures of the ligand-bound Mac1 protein. Thus, we were in the unusual position of try to get looser packing and poorer resolution. Adding a single N-terminal residue to the protein (nsp3 residues 206-374) was enough to change the packing from orthorhombic (P2_1_2_1_2_1_) to monoclinic (P2_1_). To verify binding, each of the compounds in Fig. 2 was both co-crystallized with this new Mac1 construct and soaked into crystals of the apoprotein. Despite considerable effort, we were only able to obtain a complex structure with cAMP. This model contains one molecule of Mac1 in the asymmetric unit, comprising 166 amino acids (residues 208-373), 135 water molecules, and one molecule of cAMP. The cAMP binds in the cleft between the β2-α2 loop (residues K248-V253) and the β5-α5 loop (residues L331-D339); the bottom of this cleft is formed by strand β2 (Fig. 4A-C). The electron density is well defined for all but the solvent-exposed edge of the adenine base, which appears to be wobbling at the brink of the binding site. The interactions with Mac1 are comprised entirely of hydrogen bonding interactions to the main chain, particularly in the β5-α5 loop (Fig. 4D). The adenine base is held only by water-mediated interactions to the β2-α2 and β5-α5 loops (*e*.*g*. the amide nitrogen atoms of V253 and I335) and a stacking interaction with the G251-G252 peptide bond. There are also two water-mediated contacts with a symmetry-related Mac1 molecule (Fig. 4D). Since cAMP makes no close contacts to this symmetry mate, we do not believe that the proximity of the neighboring Mac1 molecule significantly influences the binding pose of the ligand. The 2’ hydroxyl group of cAMP make hydrogen bonding interactions with the carbonyl group of A242 on strand β2 and the amide group of A254 on helix α2 (Fig. 4D). On the other side of the ribose ring, O4’ interacts with the side chain of N244 through the intercession of the water molecule. The 3’ and 5’ oxygen atoms of the ribose moiety interact, through water, with the amide of I335 and the carbonyl of A243, respectively. Given the density of interactions with the Mac1 protein, the two phosphate oxygen atoms are likely the main drivers of cAMP binding. The β5-α5 loop forms a string of amide groups that lock the phosphate of cAMP in place with hydrogen bonding interactions to these two oxygen atoms.

**Figure 4.**
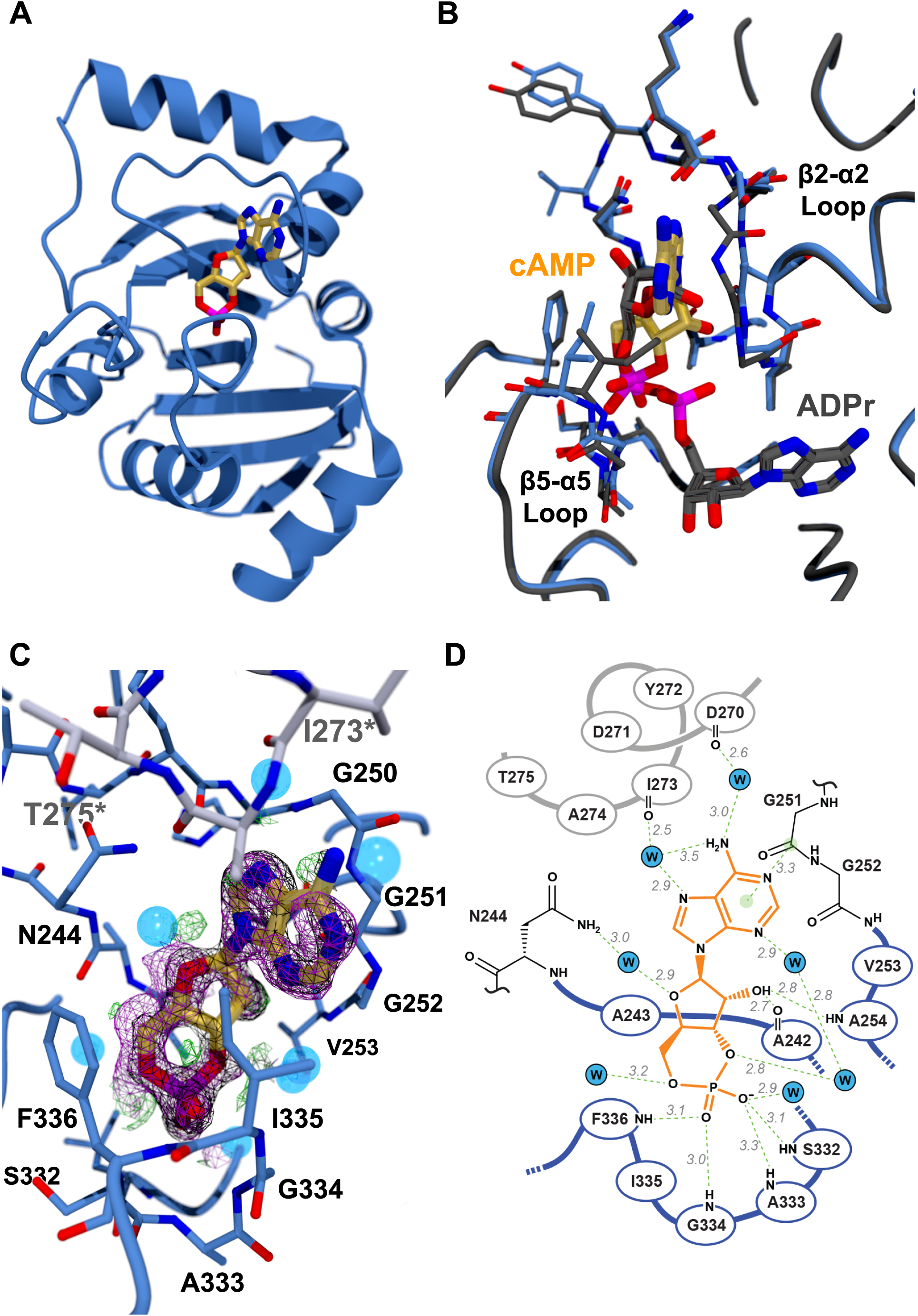
The Mac1 protein binds cAMP in an unexpected way. (A) Ribbon diagram showing of the SARS-CoV-2 MaC1 domain with cAMP bound (gold sticks). (B) Overlay of the structures of the Mac1 domain with cAMP (gold sticks) or ADPr (grey sticks) bound. The β2-α2 and β5-α5 loops are noted for reference. Note the difference in conformation of the β2-α2 loop (the section carrying G251). The reorientation of this loop allows it to pack against the adenine base of cAMP. (C) The cAMP-binding site. The Mac1 domain protein is shown as a blue Cα trace with important residues shown as thin blue sticks. Water molecules are shown as transparent blue spheres. The stretch of amino acids shown in grey sticks represents the symmetry-related Mac1 molecule that makes contacts with cAMP. The simulated annealing composite omit map is shown as a magenta mesh contoured at 1.0σ, the 2Fo-Fc map is shown in black, also at 1.0σ, and the Fo-Fc map is shown as green and red mesh at +3.0σ and -3.0σ, respectively. (D) Schematic representation of the Mac1·cAMP complex showing polar interactions between the enzyme and ligand. The cAMP ligand is shown with orange bonds. The heavy blue lines and residues drawn with black bonds represent the Mac1 protein. The heavy grey line represents the symmetry-related molecule that makes solvent-mediated contacts with cAMP. Solvent molecules are shown as blue circles with a “W”. Potential hydrogen-bonding interactions are shown as dashed green lines with the associated distances in grey italics. The stacking interaction described in the text between the G251-G252 peptide bond is shown as a green line with light green circles at each end.

The binding mode of cAMP was compared to those of ADPr (PDB ID 6W02 ^10^; Figure 4C) and AMP (PDB ID 6W6Y ^10^; not shown). Chain A of 6W02 was superimposed onto the Mac1·cAMP model using the SSM algorithm^22^ as implemented in COOT. The two models fit with a root mean square deviation (RMSD) of 0.41 Å for all 166 Cα atoms in the Mac1·cAMP model. Overlaying the AMP complex structure gave an RMSD of 0.40 Å for all Cα atoms. These low RMSD values indicate that the structures are identical in gross terms, with only small differences in the orientations of small portions such as surface loops, as one would expect when comparing multiple structures of the same protein. What is interesting is that whereas the common portions of ADPr and AMP overlay almost perfectly, cAMP binds such that the cyclic phosphate matches with the β-phosphate of ADPr, the ribose moiety corresponds to the terminal ribose of ADPr, and the adenine base is directed toward solvent. Consequently, there is no overlap at all of cAMP and AMP. This was entirely unexpected, since the adenine bases in the ADPr- and AMP-bound Mac1 domain structures are very solidly bound, with strong electron density and low B-factors. The only possible explanation for this is that the geometry of the cyclic phosphate moiety, particularly its relationship to the ribose ring, does not comport well with the α-phosphate-binding site of Mac1, and is instead a better fit for the β-phosphate/terminal ribose-binding site. This alternative binding pose results in slight reorientations of the β2-α2 and β5-α5 loops (Fig. 4C), which move away from each other to accommodate the adenine base of cAMP. It is also intriguing that, if the cAMP in this model were joined to the AMP in 6W6Y by a phosphodiester bond (and the P-O3’ bond in cAMP were broken), the result would be reminiscent of diadenosine 5’,5’-diphosphate, or related compounds like NAD(H).

## Discussion

The idea that SARS-CoV Mac1 functions as an enzyme in the cell to remove ADPr from antiviral proteins suggests that Mac1 might be an important new drug target for COVID-19.^23^ A thermal-shift binding assay was therefore developed to facilitate those efforts. The main drawback with the DSF assay was the high percentage of hits for some libraries. Various methods to successfully prioritize hits are described above, with the simplest being an examination of the initial fluorescence values in DSF assays (Fig. 3A). DSF’s other main disadvantage is that it requires relatively large amounts of protein, but this is a minor concern because of the ease with which Mac1 can be produced from *E. coli*.^3^ The most attractive alternative to a DSF binding assay would be an enzyme assay that monitors the ability of Mac1 to hydrolyze ADPr-based substrates, which are presently under development.^24^

The most intriguing hits in DSF screens were the steroids and telmisartan (Fig. 3). Unfortunately, closer examination revealed that the steroid effects in DSF appeared to be artifacts. Caution should also be exercised because telmisartan and similar compounds lowered the apparent T_m_ of SARS-CoV-2 Mac1. This could mean they bind to the protein’s unfolded state, but it is worth noting that similar destabilizing compounds were found to inhibit macrodomains in assays not based on thermal shifts,^24^ suggesting that the destabilizing compounds might bind a folded protein that assumes a different conformation.

The idea that SARS-CoV-2 ADP-ribose-binding cleft can accommodate other ligands is supported by the X-ray structure of Mac1-bound cAMP. Surprisingly the adenine base of cAMP does not bind in the adenine-binding cleft identified in the structures of Mac1 bound to ADPr or AMP.^10^ Instead, cAMP binds in the site occupied by the β-phosphate/terminal ribose unit of ADPr. This result underscores the importance of computational modelling and experimental structure determination in assessing the hits from high-throughput screening campaigns. Based on this crystal structure, it is likely that scaffolds containing a central phosphate (or diphosphate) or sulfate group could be expected to bind to this same ADPr-binding cleft. The interaction between Mac1 and cAMP, which binds Mac1 with a similar affinity as ADPr,^3^ points to other possible biological roles for Mac1, and hints that cyclic mono- or dinucleotide second messengers might allosterically modulate other nsp3 activities.

The next step in this project will be to examine the effect of promising antiviral compounds on cells harboring SAR-CoV-2 or surrogate reporter viruses. Some of the compounds above might inhibit SARS-CoV-2 replication based on the fact that Shimizu *et al*. showed that small molecules found in virtual screens targeting the homologous nsp3 macrodomain from Chikungunya virus inhibit replication of Chikungunya replicons.^25^ However, further chemical optimization would most likely be necessary for these probes to be useful in cellular studies. Fortunately, many hits reported here are already FDA approved drugs with hundreds of analogs available to facilitate such work. Alternatively, this optimized DSF assay could be used to screen larger more diverse libraries for more attractive probe candidates.

## Acknowledgements

The authors are grateful to Wilfred Tysoe and Alexander (Leggy) Arnold, for helpful discussions & technical assistance, and Daad Saffarini for allowing this work to proceed during the COVID-19 pandemic lockdown. This work was supported by University Wisconsin-Milwaukee’s generous F&A cost return policy (National Institutes of Health Grant R01 AI088001) and is based upon work supported by the National Science Foundation under Grant Number CHE-1855199 and CHE-1903899 (NV and NRS). This research used resources of the Advanced Photon Source, a U.S. Department of Energy (DOE) Office of Science User Facility operated for the DOE Office of Science by Argonne National Laboratory under Contract No. DE-AC02-06CH11357. Use of the LS-CAT Sector 21 was supported by the Michigan Economic Development Corporation and the Michigan Technology Tri-Corridor (Grant 085P1000817).

## REFERENCES

1. Shannon, A.; Le, N. T.; Selisko, B.; et al. Remdesivir and SARS-CoV-2: Structural requirements at both nsp12 RdRp and nsp14 Exonuclease active-sites. Antiviral Res. 2020, 178, 104793.

2. Freitas, B. T.; Durie, I. A.; Murray, J.; et al. Characterization and Noncovalent Inhibition of the Deubiquitinase and deISGylase Activity of SARS-CoV-2 Papain-Like Protease. ACS Infect Dis 2020, 6, 2099–2109.

3. Frick, D. N.; Virdi, R. S.; Vuksanovic, N.; et al. Molecular Basis for ADP-Ribose Binding to the Mac1 Domain of SARS-CoV-2 nsp3. Biochemistry 2020, 59, 2608–2615.

4. Ericsson, U. B.; Hallberg, B. M.; Detitta, G. T.; et al. Thermofluor-based high-throughput stability optimization of proteins for structural studies. Anal. Biochem. 2006, 357, 289–298.

5. Pantoliano, M. W.; Petrella, E. C.; Kwasnoski, J. D.; et al. High-density miniaturized thermal shift assays as a general strategy for drug discovery. J Biomol Screen 2001, 6, 429–440.

6. Huynh, K.; Partch, C. L. Analysis of protein stability and ligand interactions by thermal shift assay. Curr Protoc Protein Sci 2015, 79, 28.9.1-28.9.14.

7. Trott, O.; Olson, A. J. AutoDock Vina: improving the speed and accuracy of docking with a new scoring function, efficient optimization, and multithreading. J Comput Chem 2010, 31, 455–461.

8. Lee, P. H.; Huang, X. X.; Teh, B. T.; et al. TSA-CRAFT: A Free Software for Automatic and Robust Thermal Shift Assay Data Analysis. SLAS Discov 2019, 24, 606–612.

9. Bai, N.; Roder, H.; Dickson, A.; et al. Isothermal Analysis of ThermoFluor Data can readily provide Quantitative Binding Affinities. Sci Rep 2019, 9, 2650.

10. Michalska, K.; Kim, Y.; Jedrzejczak, R.; et al. Crystal structures of SARS-CoV-2 ADP-ribose phosphatase: from the apo form to ligand complexes IUCrJ 2020, 7,

11. Otwinowski, Z.; Minor, W. Processing of X-ray diffraction data collected in oscillation mode. Methods Enzymol. 1997, 276, 307–326.

12. McCoy, A. J.; Grosse-Kunstleve, R. W.; Adams, P. D.; et al. Phaser crystallographic software. J Appl Crystallogr 2007, 40, 658–674.

13. Emsley, P.; Lohkamp, B.; Scott, W. G.; et al. Features and development of Coot. Acta Crystallogr D Biol Crystallogr 2010, 66, 486–501.

14. Liebschner, D.; Afonine, P. V.; Baker, M. L.; et al. Macromolecular structure determination using X-rays, neutrons and electrons: recent developments in Phenix. Acta Crystallogr D Struct Biol 2019, 75, 861–877.

15. Afonine, P. V.; Grosse-Kunstleve, R. W.; Echols, N.; et al. Towards automated crystallographic structure refinement with phenix.refine. Acta Crystallogr D Biol Crystallogr 2012, 68, 352–367.

16. Cho, C. C.; Lin, M. H.; Chuang, C. Y.; et al. Macro Domain from Middle East Respiratory Syndrome Coronavirus (MERS-CoV) Is an Efficient ADP-ribose Binding Module: CRYSTAL STRUCTURE AND BIOCHEMICAL STUDIES. J. Biol. Chem. 2016, 291, 4894–4902.

17. Malet, H.; Coutard, B.; Jamal, S.; et al. The crystal structures of Chikungunya and Venezuelan equine encephalitis virus nsP3 macro domains define a conserved adenosine binding pocket. J. Virol. 2009, 83, 6534–6545.

18. Zhang; Chung; Oldenburg A Simple Statistical Parameter for Use in Evaluation and Validation of High Throughput Screening Assays. J Biomol Screen 1999, 4, 67–73.

19. Li, K.; Frankowski, K. J.; Belon, C. A.; et al. Optimization of potent hepatitis C virus NS3 helicase inhibitors isolated from the yellow dyes thioflavine S and primuline. J. Med. Chem. 2012, 55, 3319–3330.

20. Mukherjee, S.; Weiner, W. S.; Schroeder, C. E.; et al. Ebselen inhibits hepatitis C virus NS3 helicase binding to nucleic acid and prevents viral replication. ACS Chem Biol 2014, 9, 2393–2403.

21. Layton, C. J.; Hellinga, H. W. Thermodynamic analysis of ligand-induced changes in protein thermal unfolding applied to high-throughput determination of ligand affinities with extrinsic fluorescent dyes. Biochemistry 2010, 49, 10831–10841.

22. Krissinel, E.; Henrick, K. Secondary-structure matching (SSM), a new tool for fast protein structure alignment in three dimensions. Acta Crystallogr D Biol Crystallogr 2004, 60, 2256–2268.

23. Claverie, J. M. A Putative Role of de-Mono-ADP-Ribosylation of STAT1 by the SARS-CoV-2 Nsp3 Protein in the Cytokine Storm Syndrome of COVID-19. Viruses 2020, 12,

24. Wazir, S.; Maksimainen, M. M.; Alanen, H. I.; et al. Activity-Based Screening Assay for Mono-ADP-Ribosylhydrolases. SLAS Discov 2020, 2472555220928911.

25. Shimizu, J. F.; Martins, D. O. S.; McPhillie, M. J.; et al. Is the ADP ribose site of the Chikungunya virus NSP3 Macro domain a target for antiviral approaches Acta Trop 2020, 207, 105490.

