## Supplementary material for "Discovery of Drug-like Ligands for the Mac1 Domain of SARS-CoV-2 Nsp3": Table S1

**Table S1.** Crystallographic data collection and model refinement statistics for the SARS-CoV-2 Mac1 domain in complex with cAMP.

| | |
|---|---|
| **Data collection** | |
| Resolution range (Å) (last shell)[a] | 24.70 - 1.55 (1.61 - 1.55) |
| Space Group | P $2_1$ |
| *a*, *b*, *c* (Å) | 37.2, 33.0, 60.5 |
| α, β, γ (°) | 90.0, 96.4, 90.0 |
| $R_{merge}$[a] | 0.076 (0.291) |
| $R_{meas}$[a] | 0.091 (0.344) |
| $R_{pim}$[a] | 0.049 (0.182) |
| $CC_{1/2}$[a] | 0.992 (0.891) |
| No. of unique reflections[a] | 20508 (1938) |
| Completeness (%)[a] | 95.05 (90.9) |
| Multiplicity[a] | 3.4 (3.3) |
| ⟨ $I/\sigma(I)$⟩[a] | 10.9 (4.2) |
| **Model Refinement** | |
| Reflections used in refinement | 20433 (1938)[b] |
| Reflections used for $R_{free}$ | 1998 (189) |
| $R_{cryst}$ ($R_{free}$) | 0.157 (0.182) |
| Average B factor (Å$^2$) | 20.3 |
| Protein atoms | 19.1 |
| Solvent | 32.0 |
| Ligand | 17.2 |
| Root-mean-square (RMS) deviations | |
| Bond lengths (Å) | 0.008 |
| Bond angles (°) | 0.893 |
| Coordinate error (Å) | 0.11 |
| Ramachandran statistics | |
| Favored/allowed/outliers (%) | 99.4/0.6/0.0 |
| Rotamer outliers (%) | 0.8 |
| Clashscore | 0.8 |

[a]Values in parentheses apply to the high-resolution shell indicated in the resolution row

[b]The limits of the high-resolution bin for refinement were 1.59 – 1.55Å
